# Synthetic *Yarrowia lipolytica* consortium for efficient conversion of lignocellulosic oligosaccharides into lipids

**DOI:** 10.64898/2025.12.19.695487

**Authors:** Diego Crespo-Roche, Daniel González-García, Juan A. Méndez-Líter, Rodrigo Ledesma-Amaro, Alicia Prieto, Jorge Barriuso

## Abstract

Lignocellulosic biomass (LCB) is an abundant and renewable feedstock for the sustainable production of bioproducts; however, its industrial exploitation is limited by its complex composition and by the lack of microbial platforms capable of simultaneously degrading and assimilating cellulose- and hemicellulose-derived oligosaccharides. *Yarrowia lipolytica* lacks the native enzymatic machinery required for this process. In this study, we engineered a multifunctional strain (YBXT-XR-BGL3) able to secrete fungal β-glucosidase (BGL3 or BGL1) and β-xylosidase (BxTw1) from *Talaromyces amestolkiae*, enabling the hydrolysis of cellobiose and xylooligosaccharides, respectively. In addition, a xylose reductase pathway was introduced to confer xylose assimilation.

Because the construction of a single multifunctional strain may impose a significant metabolic burden and reduce fitness, we benchmarked this strain against a division-of-labor strategy. To this end, we also developed a cellobiose-specialized strain (YBGL3 or YBGL1) and a xylooligosaccharide-specialized strain (YBXT-XR). Functional characterization revealed efficient saccharification of cello- and xylooligosaccharides under acidic conditions, with BGL3 outperforming BGL1 in glucose release and BxTw1 exhibiting broad pH tolerance. Under nitrogen-limited conditions, this enabled lipid accumulation of up to 20% from cellobiose in YBGL3 and YBXT-XR-BGL3, and up to 15% from xylooligosaccharides in YBXT and YBXT-XR-BGL3.

In co-culture experiments using a mixed substrate (glucose, cellobiose, and xylooligosaccharides), both the multifunctional strain and the consortium produced up to 0.67 g L□¹ of lipids. However, the division-of-labor approach led to higher lipid accumulation (34% versus 26.3% in the monoculture), driven by a rapid population shift: following cellobiose depletion (after 72 h), YBXT-XR became predominant and utilized the remaining xylooligosaccharides almost exclusively for lipid synthesis.

Overall, this study provides the first demonstration of a *Y. lipolytica* system capable of simultaneously utilizing cellulose- and hemicellulose-derived oligosaccharides. Moreover, benchmarking a division-of-labor consortium against a multifunctional monoculture highlights a robust strategy to enhance lipid biosynthesis and improve process resilience for LCB valorization.

## 1. Introduction

Biofuel production from renewable and low-cost sources is essential to achieving sustainability and economic viability at an industrial scale. In this context, lignocellulosic biomass (LCB) stands out as a promising feedstock to produce high-value-added products due to its abundance and availability ^1^. Furthermore, the expansion of global agricultural production has increased the generation of agro-industrial residues, positioning LCB as a critical resource in the frame of bioeconomy. Valorization of these residues supports the transition to a circular economy, where waste from one process is used as feedstock for another ^2^, and where LCB biorefineries replace crude oil-based refineries ^3^. However, the intricate structure of LCB poses significant challenges that must be addressed for efficient conversion into fermentable sugars and subsequent production of the targeted products. Usually, LCB pretreatment is needed to make cellulose and hemicellulose accessible for the enzymes ^4^. Typically, acid or alkaline conditions are used, in some conditions these pretreatments release celo- and xylooligosaccharides ^5^ that are potentially valorizable by engineered strains.

LCB is primary composed of lignin, a highly recalcitrant aromatic polymer formed by phenylpropanoid units mainly linked by β-O-4 and C–C bonds, and two polysaccharides: cellulose and hemicellulose. Cellulose, the most abundant, is a linear homopolymer composed of D-glucopyranose subunits linked by β-1,4-glycosidic bonds, while hemicelluloses comprise a group of branched heteropolymers. In most cases, their backbone consists of β-(1→4)-linked sugars, with side chains made up of different monosaccharides. Xylans, which are among the most abundant hemicelluloses, consist mainly of a β-1,4-linked xylose backbone that may be substituted with arabinose, glucuronic acid, and acetyl groups ^6^. Cellulose and xylan depolymerization release D-glucose and mainly D-xylose respectively.

In nature, wood-degrading fungi are the main responsible of LCB degradation through the secretion a broad array of enzymes ^7^. The ascomycete *Talaromyces amestolkiae*, is known for releasing a wide enzymatic arsenal for LCB degradation ^8,9^. Among the extracellular cellulolytic enzymes characterized from this filamentous fungus, the β-glucosidases BGL1 ^10^ and BGL3 ^11^ are able to degrade cellobiose and longer cellooligosaccharides (β-1,4-D-glucopyranose oligomers) to glucose. Similarly, the extracellular β-xylosidase BxTw1 from this fungus hydrolyzes xylooligosaccharides containing 2 to 6 xylose units into xylose, and exhibit one of the highest catalytic efficiencies reported in the literature ^12^. These three enzymes catalyze the final steps of lignocellulosic biomass degradation, releasing metabolizable sugar monomers.

*Yarrowia lipolytica* is a dimorphic yeast with high industrial interest due to its robustness and stability during fed-batch fermentations ^13^, and its ability to accumulate triacylglycerides in lipid bodies under nitrogen- or phosphorus-limiting conditions ^14^. This characteristic is enhanced by an efficient acetyl-CoA ^15^ metabolic pathway and pentose phosphate pathway (PPP) ^16^ that provide with precursors and reducing power respectively, for the production of lipids. Microbial lipid production has attracted considerable interest as a sustainable route to produce biofuels (especially for aviation) and bulk chemicals from low-cost carbon feedstocks ^17–20^. In this context, *Y. lipolytica* engineered strains with a substantially enhanced capacity for lipid accumulation have been developed through a combination of metabolic rewiring strategies, including reinforcement of the native fatty-acid biosynthetic pathway, attenuation of competitive β-oxidation, optimization of acetyl-CoA and NADPH supply, and manipulation of key regulators of lipid storage ^21–23^. These engineering efforts have yielded strains capable of accumulating lipids at significantly higher titers and productivities than their wild-type counterparts. However, *Y. lipolytica* cannot grow on LCB as it lacks the enzyme repertoire required for its depolymerization Additionally, this yeast can metabolize glucose but not xylose, the second most abundant sugar released from LCB ^24^. Therefore, several strategies have been used to address these issues. For example, activation of the cryptic xylose metabolism ^25,26^ or expression of xylose isomerase and/or xylose reductase pathways ^27^ allowing xylose assimilation. Other approaches aimed at exploiting cellulose and xylan from LCB have focused on the heterologous expression of fungal cellulases and hemicellulases in *Yarrowia* ^28–30^. Lipids production from cellulosic feedstocks has been studied over the last decade, achieving up to 5 g L^-1^ of lipids from glucose and cellulose in a bioreactor using a single *Y. lipolytica* strain expressing several fungal cellulolytic enzymes ^31^. Other value-added chemicals, such as citric acid, have also been produced from these feedstocks ^32^.

To date, no *Y. lipolytica* strain capable of simultaneously degrading cellulose and hemicellulose and metabolizing the resulting products has been reported. Probably, endowing a single *Y. lipolytica* chassis with all the enzymatic activities required would impose a substantial metabolic burden ^33,34^, potentially reducing cellular performance and limiting its industrial applicability. However, this would enable the concurrent valorization of the two major lignocellulosic polysaccharides, improving substrate flexibility and industrial viability of microbial lipid production ^35,36^. The use of microbial consortia, leveraging division of labor, emerges as a compelling alternative strategy, in which different strains are engineered to specialize in different tasks, such as polymer depolymerization, sugar assimilation, or lipid accumulation. In this setup, nutritional demands can be distributed across the populations in the consortium, alleviating the metabolic load comparted to the engineering of a single chassis ^37^. The challenge of the establishment of stable yeast consortia and division of labor has been studied in the model organism *Saccharomyces cerevisiae* ^38,39^, demonstrating improved bioproduction yields of flavonoids ^40,41^ and ethanol from plant biomass ^42^. Recent examples of *Y. lipolytica*-based consortia for revalorization of LCB have also been developed, in combination with immobilized *Trichoderma reesei* ^43^, *Cellulomonas fimi* ^44^ and a co-culture of three *Y. lipolytica* strains expressing three different fungal cellulolytic enzymes ^29^. However, the yields in the latter were poor (188 mg L^-1^ of lipids), probably due to the lack of sufficient β-glucosidase activity in the culture and suboptimal inoculation ratios for maximizing lipid production.

In this work, we focused on developing degrading-specialized and nutritionally-codependent *Y. lipolytica* strains for lipid production from cello- and xylo-oligosaccharides, which were subsequently evaluated in co-culture. These recombinant strains contained the *bgl3* and *bxtw1* genes from *T. amstolkiae* and the xylose reductase metabolic pathway from *Scheffersomyces stipites* (with xylulokinase from *Y. lipolytica*). This division-of-labor strategy was assessed to determine whether the cellular stress of each specialized strain was kept at levels compatible with high process efficiency, benchmarking against a monoculture engineered to carry all the functionalities within a single chassis. This study provides the first demonstration of a *Y. lipolytica* system capable of simultaneously utilizing both cellulose and hemicellulose-derived oligosaccharides, ultimately enhancing lipid biosynthesis and strengthening process robustness.

## 2. Materials and methods

### 2.1. Strains and culture media

The microorganisms used and designed in this study and their genetic background are summarized in Table 1. *Y. lipolytica* W29 was genetically modified for increased lipids accumulation, obtaining the *Y. lipolytica* DGA ^27^ strains (YDGA or YDGA Leu^-^ for the leucine auxotroph). *E. coli* DH5α was used for plasmid construction and maintenance.

**Table 1.** Plasmids and strains used in this study.

| Plasmid | Description | Source of reference |
| --- | --- | --- |
| JMP62Leu | pTEF, LEU2 | 45 |
| JMP62Ura | pTEF, URA3 | 45 |
| JMP62Leu-xylred | D-xylose reductase from *Scheffersomyces stipites* (P31867), xylitol dehydrogenase from *S. stipites* (E5G6H4), xylulokinase from *Y. lipolytica* W29 (YALI0F10923), LEU2 | 27 |
| JMP62-BGL1 | BGL1 with SP3 signal peptide under pTEF control, LEU2 | This work |
| JMP62-BGL3 | BGL3 with SP3 signal peptide under pTEF control, LEU2 | This work |
| JMP62-BXT | BxTw1 with SP3 signal peptide under pTEF control, LEU2 | This work |

| Strain | Relevant genotype | Source of reference |
| --- | --- | --- |
| *Escherichia coli* |  |  |
| DH5α | F<sup>-</sup> ϕ80lacZAM15 $\Delta$(lacZYA-argF)U169 recA1 endA1 hsdR17(rK<sup>-</sup>, mK<sup>+</sup>) phoA supE44 λ<sup>-</sup> thi<sup>-1</sup> gyrA96 relA | Thermo Fisher Scientific |
| *Yarrowia lipolytica* |  |  |
| Pod1 | W29 ura3-302 leu2-270 xpr2-322 (Leu<sup>-</sup>, Ura<sup>-</sup>, ΔAEP, Suc<sup>+</sup>) | 49 |
| YDGA Leu<sup>-</sup> | Po1d with *dga1* gene from *Rhodospiridium toruloides* and *dga2* gene from *Claviceps purpurea* inserted in *mef1* gene | CIB collection |
| YDGA | YDGA transformed with plasmid JMP62Leu | This work |
| YBGL1 | YDGA transformed with plasmid JMP62-BGL1 | This work |
| YBGL3 | YDGA transformed with plasmid JMP62-BGL3 | This work |
| YBXT | YDGA transformed with plasmid JMP62-BXT | This work |
| YBXT-XR | YBXT transformed with plasmid JMP62-xylred | This work |
| YBXT-XR Leu<sup>-</sup> Ura<sup>-</sup> | YBXT-XR with leucine and uracil auxotrophy | This work |
| YBXT-XR-BGL3 | YBXT-XR Leu<sup>-</sup> Ura<sup>-</sup> transformed with JMP62-BGL3 and JMP62Ura | This work |

*E. coli* was cultured in Luria-Bertani medium (LB, 10 g L^-1^ NaCl, Fischer Scientific; 10 g L^-1^ bactotryptone, Gibco; and 5 g L^-1^ yeast extract, Gibco) at 37 °C and 180 rpm. Biomass evolution was followed by measuring the optical density (OD) of the cultures at 600 nm in a Shimadzu UV-260 spectrophotometer. *Y. lipolytica* strains were cultured in the rich medium YPD (10 g L^-1^ yeast extract, 20 g L^-1^ bactopeptone, Condalab; and 20 g L^-1^ glucose, VWR) and the minimum medium YNB (Difco, 6.7 g L^-1^ YNB salts), supplemented with leucine (Sigma Aldrich, Drop-out Medium Supplements without Uracile) when required at 30 °C and 180 rpm. Glucose (VWR), xylose (Thermo Scientific), cellobiose (Sigma), xylooligosaccharides (Anhui Elite Industrial CO, C2-C5) or a mixture of them, were used as carbon sources.

Transformants were selected on YNB agar plates with glucose or xylose (10 g L^-1^) without leucine (1.5 % w/v agar, pH 7). To determine enzymatic activities, the strains were cultured in YNB with 20 g L^-1^ of glucose at 30 °C and 180 rpm.

Different inoculum ratios between the two recombinant strains YBXT-XR (xylooligosaccharides degrader) and YBGL3 (cellobiose degrader) were assayed in experiments with consortia. The initial OD_600nm_ in the co-cultures was always 0.3, ensuring that the contribution of each strain was at least 0.025. The ratios assayed were 1:1, 1:2, 1:5 and 1:10, always in favor of the YBXT-XR strain, as xylooligosaccharides degradation was the limiting step in these assays. At these ratios, the OD_600nm_ YBXT-XR/YBGL3 were 0.15/0.15, 0.10/0.20, 0.05/0.25, 0.025/0.250.

### 2.2. Plasmid construction and transformation of *Y. lipolytica* DGA strains

The plasmids employed in this study are summarized in Table 1. All genes have been designed with a upstream constitutive pTEF promoter, a XPR2 downstream terminator in the plasmid pJMP62 ^45^ and the secretory signal peptide SP3 ^46^. The genes coding for β-glucosidase 1 (BGL1) ^10^, β-glucosidase 3 (BGL3) ^11^ and β-xylosidase (BxTw1) ^12^ of *T. amestolkiae* were amplified by PCR with the error-proof GLX polymerase (R050A, Takara) using a set of primers (Table S1) containing the restriction site *Bam*HI and *Avr*II in the forward and reverse primer, respectively. Amplicons were cleaned with the PCR purification kit QIAquick (QIAGEN) and digested with *Bam*HI and *Avr*II for 2 hours at 37 °C. The plasmid JMP62Leu was also digested in the same conditions. The digestions were subsequently cleaned-up using the QIAquick kit and ligation was performed with T4 ligase for 15 min at RT, generating the plasmids JMP62-BGL1, JMP62-BGL3 and JMP62-BXT, respectively. Immediately, they were transformed into *E. coli* DH5α competent cells by thermal shock. Transformants were plated in LB with 1.5% agar and kanamycin (50 µg mL^-1^) as the selection antibiotic. Presence of the plasmid was checked by colony PCR with NZYTaq II 2x Green Master Mix (NZYtech) using primers F_JMP62_seq1 and R_BGL1_seq2, R_BGL3_seq3 or R_BxTw1_seq3 depending on the transformation. For subsequent plasmid extraction, PCR-positive colonies were cultured in LB medium with kanamycin, and plasmids were extracted using NZY Miniprep kit (NZYTech). Finally, the plasmids were transformed in *Y. lipolytica* by the lithium acetate method, integrating the DNA cassettes randomly in the genome of the yeast ^47^. Transformants were selected on YNB with glucose 20 g L^-1^ as the carbon source and no supplementation, as the leucine auxotrophy is reverted if the cell is transformed. The strains generated were named YBGL1, YBGL3, YBXT respectively.

The plasmid JMP62 harbouring the reductase pathway for the metabolism of xylose ^27^ was transformed into YBXT by the lithium acetate method. Selection was performed in YNB plates with 20 g L^-1^ xylose and the strain was named YBXT-XR. This strain prototrophic strain was then transformed with the plasmid harbouring the Cre recombinase to revert to an auxotrophic phenotype for further engineering. Cre recombinase recognizes the *loxP* and *loxS* sequences flanking the genes *leu2* and *ura3*, excising the DNA between them ^48^ and restoring uracil and leucine auxotrophy in YBXT-XR. Then, transformation with JMP62-BGL3 and JMP62Ura vector resulted in the generation of YBXT-XR-BGL3 prototroph strain.

### 2.3. Measurement of enzymatic activity

After *Y. lipolytica* DGA transformation, 35 positive clones of each construction were streaked in YNB plates and cultured 24 h at 30 °C for activity screening. The plates were covered with a solution of the fluorogenic substrates 50 mM 4-methylumbelliferyl-β-D-glucopyranoside (MUG) for BGL1 and BGL3 transformants, or 50 mM 4-methylumbelliferyl-β-D-xylopyranoside (MUX) for BxTw1 recombinant strains, in 100 mM sodium acetate buffer and 0.8% agar. After solidifying, the plates were placed in an oven at 50 °C for 20 min, and positive clones were revealed in a transilluminator BioRad Gel Doc XR. Clones with β-glucosidase/β-xylosidase activity hydrolyzed MUG/MUX, quantitatively releasing fluorogenic methylumbelliferone ^50^.

The β-glucosidase and/or β-xylosidase activity of the culture supernatants was measured in a heating block at 50 °C and 1000 rpm, using 0.1 % (w/v) *p*-nitrophenyl-β-D-glucopyranoside (*p*NPG, Sigma) or *p*-nitrophenyl-β-D-xylopyranoside (*p*NPX, Sigma) in sodium acetate buffer 100 mM, pH 4.0. The reaction was stopped after 10 min by adding 2 % of Na_2_CO_3_ (final concentration of 1.4 %), and the release of *p*-nitrophenol (*p*NP) was measured in a spectrophotometer at 410 nm. One activity unit of β-glucosidase was defined as the amount of enzyme capable of releasing 1 μmol of *p*NP per min (the molar extinction coefficient, ε, of *p*NP is 15,200 M^−1^ cm^−1^).

### 2.4. Supernatant concentration and enzymatic saccharification

*Y. lipolytica* cultures in YNB-glucose medium were centrifuged (10 min, 4000 rcf), after 20 h of growth and the supernatant recovered and concentrated in 10-kDa Amicon centrifugal filters (Sigma Aldrich). Enzymatic activities were monitored during the concentration process. Oligosaccharide saccharifications (10 g L^-1^ cellobiose, 1 g L^-1^ xylobiose or 10 g L^-1^ xylooligosaccharides) were carried out using 100 µL of the concentrated supernatants in a thermoblock (Thermo-Shaker, Biosan). The reactions were conducted at 1000 rpm for 18 h, at 30 or 50 °C and different pH values (5, 6 and 7). Saccharification performance (%) was calculated as the proportion of monosaccharide (glucose or xylose) relative to the total sugar concentration (glucose + cellobiose, or xylose + xylobiose + higher oligomers) and normalized to the control, as the xylooligosaccharides substrate contains a minimal quantity of monomeric xylose.

### 2.5. Yeast populations quantification

The proportion of YBL3 (only consumes glucose) and YBX-XR (consumes glucose and xylose) cells in the consortia formed by these two strains was followed by drop-plate assay. A sample of the culture was serially diluted from tenfold up to a dilution of 1:10^5^ and a drop of 10 µL of each dilution was spotted onto agar plates (YNB agar plates with 20 g L^-1^ of glucose or xylose) for colony forming units counting after 48 h of growth at 30 °C. Both YBGL3 and YBXT-XR grow on glucose while only YBXT-XR can grow on xylose, allowing the quantification of total cells in the culture (colonies observed in YNB glucose plates) and quantification of YBXT-XR cells (colonies observed in YNB xylose plates). Finally, the proportion of YBXT-XR cells over the total population was calculated.

### 2.6. HPLC analysis of sugars

Sugar concentration in the reaction mixtures was determined by High-Performance Liquid Chromatography (HPLC) using an Agilent Technologies 1200 series instrument, equipped with an Aminex HPX-42C column, 300 x 7.8 mm (BioRad) maintained at 80°C and a refractive index detector (RID). A mobile phase consisting of MilliQ water was employed at a flow rate of 0.4 mL min^−1^. For calibration, standards of glucose, xylose, cellobiose, xylobiose, xylotriose and xylopentaose at different concentrations were analyzed in the same running conditions.

### 2.7. Lipid production, visualization and measurement

YNB without ammonium sulfate and amino acids (YNBww; Thermo Scientific) was used as the basal medium for lipid production assays. Cultures were supplied with 20 g L□¹ of the corresponding carbon sources and ammonium sulfate ((NH□)□SO□ Sigma-Aldrich) supplemented as the nitrogen source to establish two distinct C/N regimes: a non-limited condition (C/N = 5; 7.5 g L□¹ (NH□)□SO□) and a nitrogen-limited condition (C/N = 75; 0.5 g L^-1^ (NH□)□SO□). Lipid accumulation was visualized by fluorescence microscopy (Leica DM4 M) after staining with BODIPY (Thermo Fischer), a fluorophore with excitation and emission peaks at 493 and 504 nm, respectively. The sample solution (10 μL) was added to the centre of the glass slide, followed by a cover glass (20 × 20 mm) sealing, and observed at 1000X magnification. For lipid extraction and quantification, culture pellets were freeze-dried and around 200 mg of dried biomass were mixed with 2 M HCl and incubated at 80 °C for 1 h. When cooled, 2 mL of methanol and 1 mL of chloroform were added and the sample subjected to vigorous vortexing. 1.8 mL of MilliQ water was then added and samples were centrifugated at 2,500 g for 15 min at 4 °C. The organic phase at the bottom was separated in a pre-weighed tube, and the aqueous phase was re-extracted with 2 mL of a 1:10 methanol/chloroform mixture. After vortexing and centrifugation, the organic phase was recovered and added to the tube containing the first organic extract. Chloroform was evaporated and the dry lipid content was weighted to calculate its percentage relative to dry biomass (using the weight of the pellet prior extraction). Total dry biomass in the culture was also measured previous to lipid extraction to calculate final lipid concentration (g L^-1^).

## 3. Results

### 3.1. The modified *Y. lipolytica* DGA strains secrete β-glucosidase and β-xylosidase activities

*Y. lipolytica* strains YBGL1, YBGL3 and YBXT were generated by transformation of the obese YDGA Leu^-^ strain with the integrative plasmids JMP62-BGL1, JMP62-BGL3 and JMP62-BXT, respectively. Then, the β-glucosidase and β-xylosidase activities of 35 candidates were assessed on solid medium using MUG and MUX assays (Fig. S1). As expected for constructs integrated through zeta elements, transformants displayed a wide range of expression levels ^51^. For each enzyme, four clones displaying distinct fluorescence intensities (named High1, High2, Medium and Low, depending on the observed fluorescence) were selected for further characterization.

Growth and extracellular enzymatic activity of the candidates were measured over 48 h in YNB with 20 g L^-1^ glucose. All selected candidates showed extracellular enzymatic activity as SP3 signal peptide enabled the efficient secretion of the enzymes, as previously reported ^46,52^. No substantial growth differences (OD_600nm_) were observed between the candidates and the control strain YDGA, except for clone High1 from YBGL3, which showed a slight growth impairment at 30 h. In terms of enzymatic activity, YDGA did not have β-glucosidase or β-xylosidase extracellular activity as expected, while the transformants presented different levels of activity depending on the expressed enzyme (Fig. S2). At 24 h, the activity in the clone Medium of BGL1 peaked at around 150 mU mL^-1^ and in BGL3 clone High2 raised up to 250 mU mL^-1^, suggesting that the β-glucosidase BGL3 outperforms BGL1 in *Y. lipolytica*. It is interesting to note that β-glucosidase activity, in both cases, was almost completely lost when the culture reached the stationary phase at 30 h. This drop can be attributed to proteolysis by the extracellular acid protease Axp, whose expression is induced upon acidification of the medium during sugar metabolism ^53^ In comparison, YBXT clones displayed lower enzymatic activities (around 30 mU mL^-1^ of β-xylosidase activity for clones Medium and Low). However, this activity did not drop and even increases in some cases in the stationary phase, suggesting higher resistance to Axp-mediated degradation—a relevant trait given the acidic characteristic of LCB hydrolysates ^5^. Based on these results, the best-performing clone was selected for each enzyme: clone Low for BGL1, clone High2 for BGL3, and clone Low for BXT, hereafter designated as YBGL1, YBGL3, and YBXT, respectively. This highlights that fluorogenic plate assays (Fig. S1) provided a first qualitative discrimination between functional and non-functional clones but liquid assays (Fig. S2) are required to identify genuinely high-activity clones.

The supernatants of 24 h-cultures of YBGL1, YBGL3 and YBXT in YNB-glucose were concentrated through 10-kDa filters (Table S2) to assay cellobiose and xylobiose saccharification. These fungal glycosyl hydrolases present acidic optimal pHs ^10–12^ while Y. lipolytica is known to decreases the medium pH when growing on sugars so a neutral pH at beginning of the culture may be suitable to delay high acidification ^54^. In this way, saccharifications were performed between pH 5 and 7 evaluate the balance between these two key aspects. As shown in Figure 1, YBGL3 and YBXT degraded cellobiose and xylobiose, respectively, in almost all conditions tested, with better performance at high temperature (50 °C) and in an acidic medium (pH 5). However, BGL1 did not release glucose in these conditions, except for a reduced saccharification at pH 7. Therefore, YBGL1 was discarded for further experiments, as the β-glucosidase BGL3 from *T. amestolkiae* outperformed BGL1 in enzymatic activity and substrate degradation.

**Figure 1.**
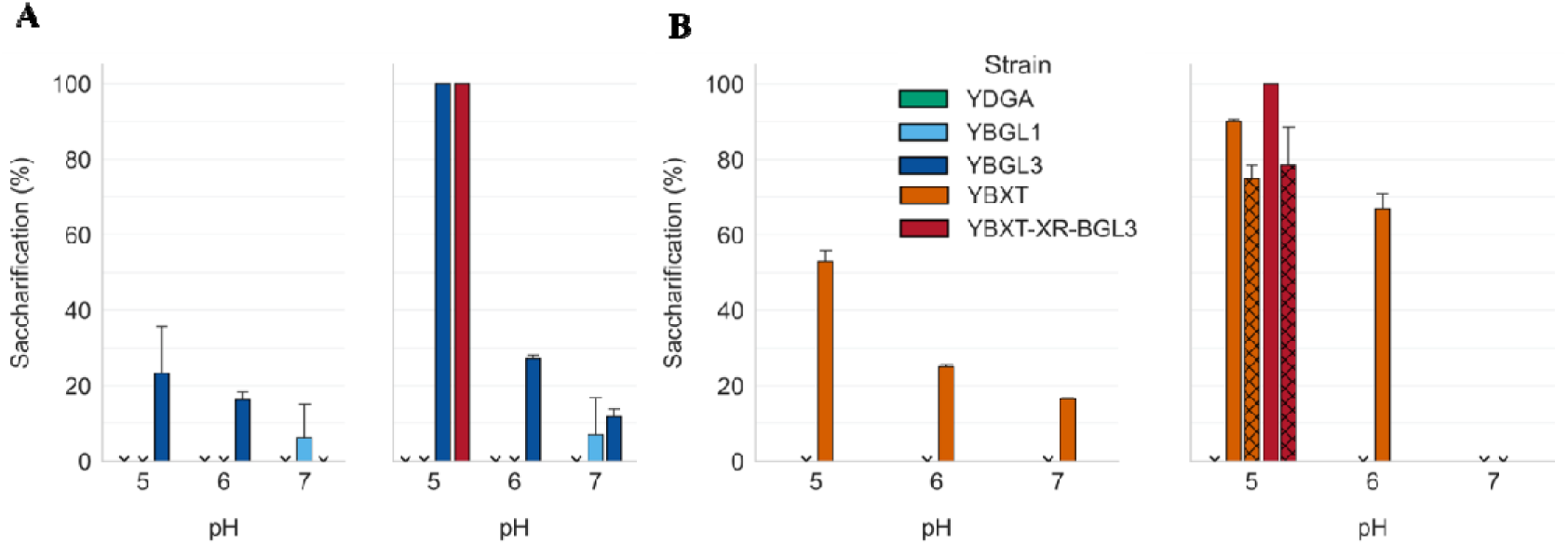
Saccharification performance of the concentrated supernatant (10X concentration) in different reactions conditions. **A)** 10 g L^-1^ cellobiose degradation at 30 °C (left side) or 50 °C (right side) at pH 5, 6 or 7. Strains tested were YDGA (green), YBGL1 (light blue), YBGL3 (dark blue) and YBXT-XR-BGL3 (red). **B)** 2 g L^-1^ xylobiose (plain bar) and xylooligosaccharides (hatched bar) degradation at 30 °C (left side) or 50 °C (right side) at pH 5, 6 or 7. Strains tested were YDGA (green), YBXT (orange) and YBXT-XR-BGL3 (red). Error bars correspond to the standard variation of technical duplicates.

Therefore, we evaluated the growth and substrate consumption profile of YBGL3 (Fig. 2A) by incubating this strain in YNB medium with 10 g L^-1^ of cellobiose supplemented with 1 g L^-1^ of glucose (to help with initial growth and protein secretion). Growth on cellobiose was compared to that of YBGL3 and the control strain YDGA in YNB medium with either 11 g L^-1^ or 1 g L^-1^ of glucose. Both strains showed similar growth over glucose at all pHs tested. On the contrary, only YBGL3 grew in cellobiose at pH 5 and 6, this growth was similar to that of YDGA and YBGL3 with 11 g L^-1^ of glucose, demonstrating the high efficiency of the enzyme BGL3 at acidic pH, where cellobiose is rapidly degraded and glucose metabolized by the yeast. However, YBGL3 was not able to use cellobiose at pH 7 as its growth was similar to YNB glucose 1 g L^-1^, where cellobiose is not present due to inefficient cellobiose hydrolysis (Fig. 2A) as BGL3 performance drops at neutral pH ^11^ but the enzyme was being produced by YBGL3 as β-glucosidase activity was detectable in supernatants assayed at pH 4. YDGA showed similar growth with 1 g L^-1^ of glucose, supplemented or not with cellobiose 10 g L^-1^, proving that it cannot degrade cellobiose.

**Figure 2.**
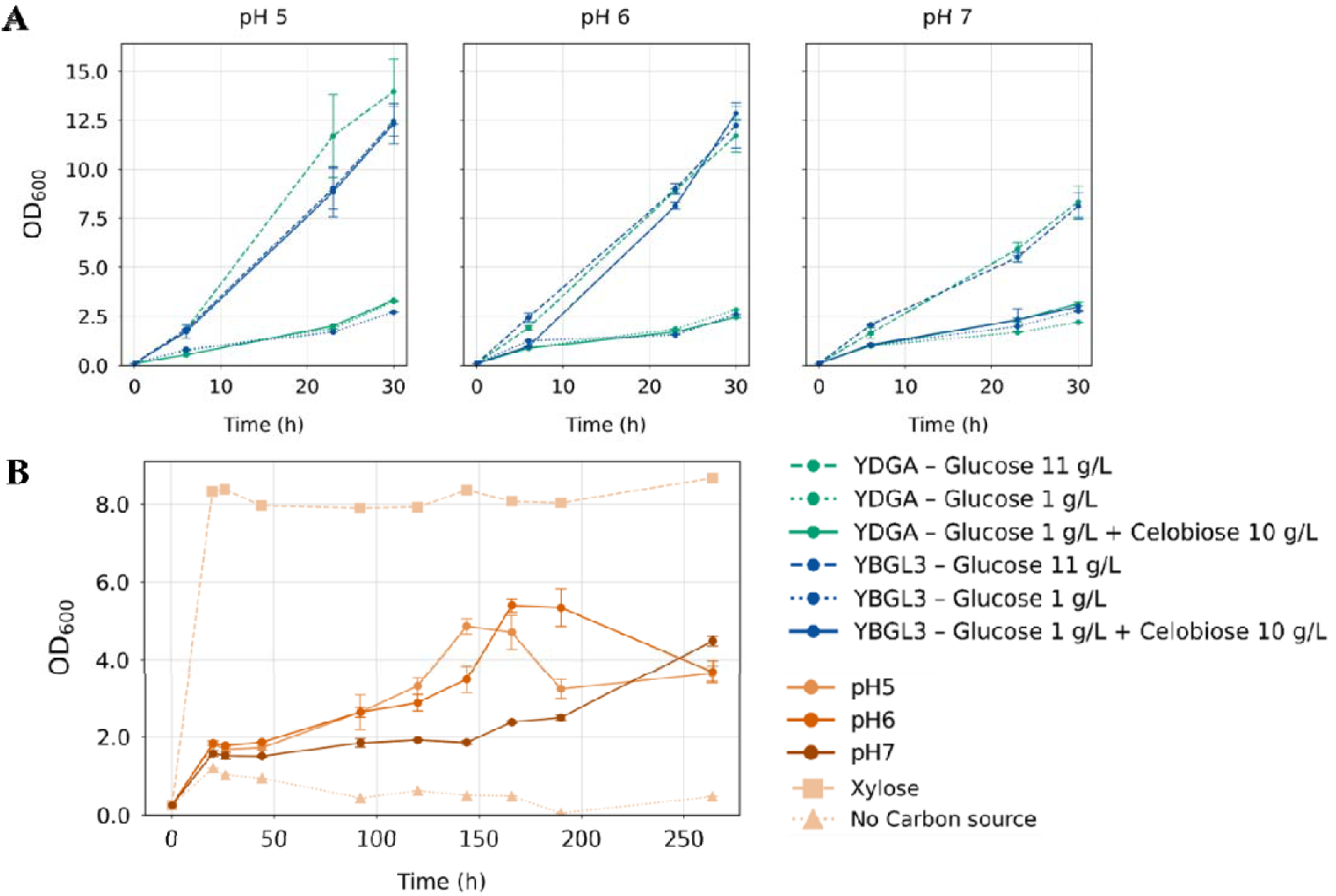
A) Growth of YDGA (green) and YBGL3 (blue) over glucose 11 g L^-1^ (dashed line), glucose 1 g L^-1^ (dotted line) or cellobiose (10 g L^-1^ and 1 g L^-1^ of glucose, solid line) at pH 5, 6 and 7. B) Growth of YBXT-XR over xylose (10 g L^-1^, dashed line) and xylooligosaccharides (10 g L^-1^ and 1 g L^-1^ xylose, solid line) at different pHs and no carbon source (dotted line). Error bars correspond to the standard variation of biological duplicates.

### 3.2. Enabling xylooligosaccharide degradation and assimilation in *Y. lipolytica* DGA

To generate a strain able to consume the xylose liberated by the recombinant β-xylosidase, YBXT was transformed with JMP62Leu harboring the xylose reductase pathway (JMP62-XR). This strain, named YBXT-XR, maintains the ability to metabolize glucose. Transformants with detected extracellular *p*NPX activity were tested against xylooligosaccharides at pH 6 and clone 1 showed the best performance, hence, it was selected for further assays (Fig. S3).

The next step was to assess whether the doubly engineered strain was capable of growing on, and metabolizing, xylose. In this case, YBXT-XR (Fig. 2B) was incubated in YNB medium with 10 g L^-1^ of xylooligosaccharides supplemented with 1 g L^-1^ of xylose at pH values of 5, 6 and 7. Growth was compared to that of the strain YBXT-XR cultured in YNB medium with 11 g L^-1^ of xylose or without carbon source.

The strain YBXT-XR showed consistent growth as a consequence of degradation and metabolization of xylooligosaccharides at pH 5, 6 and 7 (Fig. 2B), displaying substantially greater functional tolerance to pH elevation than YBL3. This behaviour is consistent with the broader pH activity profile reported for BxTw1, which is active from pH 3 to pH 7 and retaining significant activity even under mildly alkaline conditions ^12^. All growth curves reflected a rapid OD_600nm_ increase during the first 20 h of culture, with xylose being rapidly consumed, followed by a lag-phase. The control culture grown on xylose quickly entered the stationary phase and reached the highest OD_600nm_ among all tested conditions, as this substrate is directly internalized and metabolised. No major differences were observed between growth at pH 5 and 6, with a shorter lag phase and higher OD_600nm_ and β-xylosidase activity than at pH 7, which is consistent with the higher catalytic efficiency of BxTw1 at acidic pH ^12^. At pH 7, growth started again at 144 h after a long lag-phase, during which β-xylosidase activity slowly rose until 10 mU mL^-1^. Then, a great increase that correlated with a rapid yeast growth was detected at 264 h, reaching similar final OD_600nm_ than at pH 5 and 6.

YBXT-XR is able to metabolize xylooligosaccharides, however it grows slower than YBGL3 on cellobiose, this can be attributed to a lower β-xylosidase secretion in *Y. lipolytica* (Fig. S2), suggesting that glycosidase load or folding efficiency differs between the two enzymes. Second, catabolism of xylose via the XR/XDH pathway often leads to cofactor imbalance (NADPH/NADH) and xylitol overflow, which reduces metabolic efficiency ^18^.

### 3.3. Engineering a multifunctional strain for lipid production from cello- and xylooligosaccharides

Once each of the individual strains were fully characterized, we proceeded to combine all their capabilities in a single *Y. lipolytica* variant harbouring BGL3, BxTw1 and the xylose reductase pathway. The new strain, denominated YBXT-XR-BGL3, was expected to experience some metabolic cost due to the expression of two extracellular enzymes and the xylose cassette, compared to YBGL3 and YBXT-XR. However, its growth on cellobiose was similar to YBGL3 (Fig. 3A) and to YBXT-XR on xylooligosaccharides (Fig. 3B), respectively.

**Figure 3.**
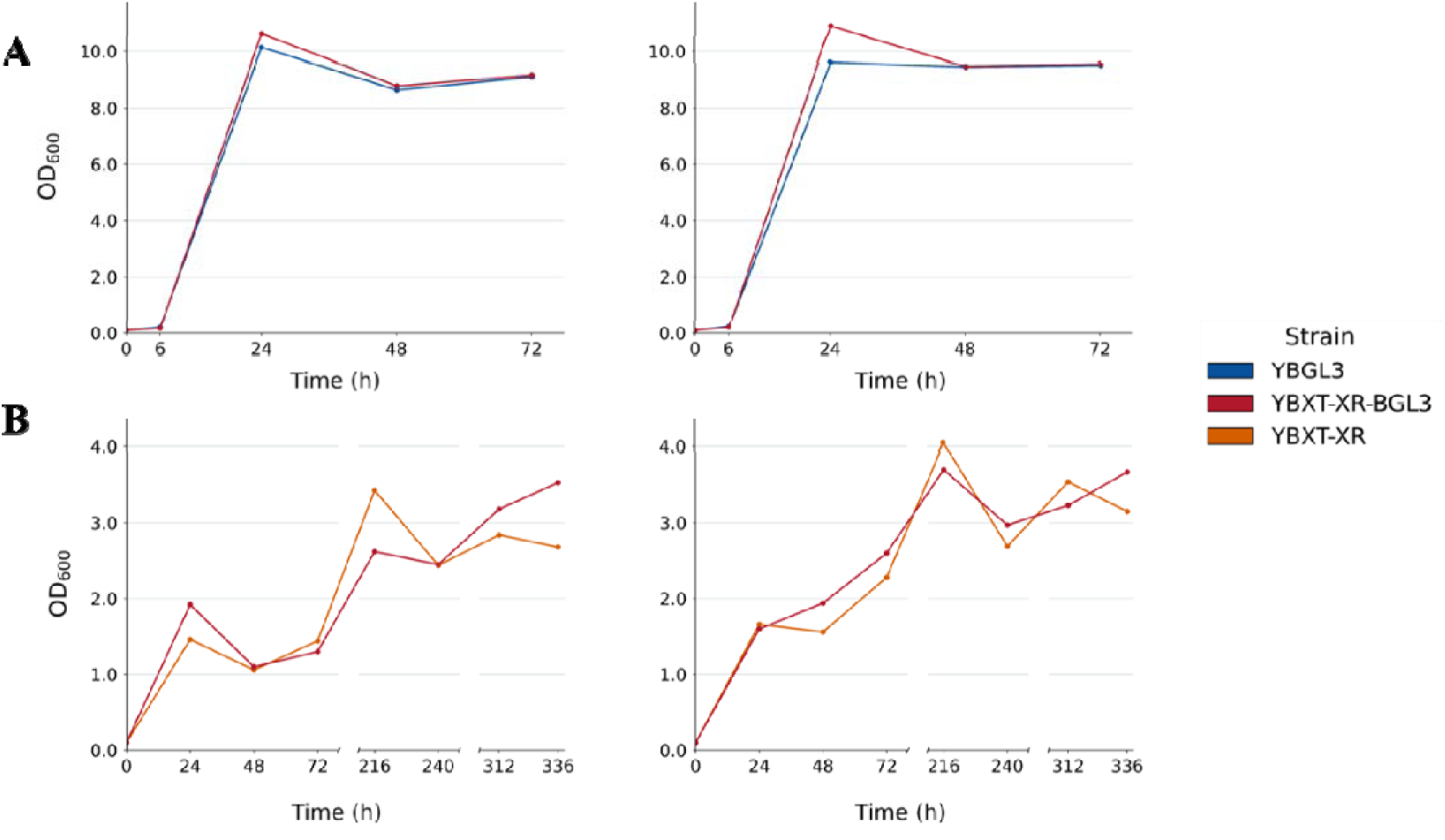
**A)** Compared growth of YBGL3 (blue) and YBXT-XR-BGL3 (red) at pH 5 (left) and 6 (right) on cellobiose 10 g L^-1^ and glucose 1 g L^-1^. **B)** Compared growth of YBXT-XR (orange) and YBXT-XR-BGL3 (red) at pH 5 (left) and 6 (right) on xylooligosaccharides 10 g L^-1^ and xylose 1 g L^-1^.

Saccharification of of cellobiose (10 g L^-1^), xylobiose (2 g L^-1^) and xylooligosaccharides (10 g L^-1^) was tested in the best conditions previously defined for YBGL3 and YBXT (Fig. 1, pH 5 and 50 °C). The concentrated supernatants of YBGL3-BXT-XR (Table S2) degraded cellobiose and xylobiose into the corresponding monosaccharide in 18 h, like those from YBGL3 (100 % saccharification) and YBXT (∼90 % saccharification) (Fig. 1). However, the saccharification of longer xylooligosaccharides, measured in the same conditions, was not complete using YBXT-XR-BGL3 and YBXT supernatants (75 % and 71 %, respectively).

### 3.4. Lipid production from lignocellulosic oligosaccharides

As a final characterization step of the microbial chassis developed in this work, their ability to produce intracellular lipids at pH 6 was evaluated. *Y. lipolytica* accumulates TAGs in lipid bodies in the cytoplasm under nitrogen-limiting conditions with an available carbon source. Thus, C/N molar ratios were adjusted to 5 and 75 in the culture media to induce lipid synthesis and accumulation. For YBGL3, the substrate consisted on cellobiose (19 g L^-1^) with 1 g L^-1^ glucose to induce initial culture growth and protein secretion, while cultures with 21 g L^-1^ of glucose served as control to provide the same molar carbon input. The strain YBXT-XR was evaluated under the same conditions but using as substrate a mixture of xylooligosaccharides (18 g L^-1^) with xylose (2 g L^-1^) or the equivalent carbon amount of xylose (21 g L^-1^). Lipid production was quantified gravimetrically once the cultures entered the stationary phase and showed lipid accumulation (Fig. S4), which was confirmed by observing lipid bodies under the microscope after BODIPY staining. The stationary phase was reached after approximately 3 days for YBGL3 cultures on glucose andcellobiose, 6 days for YBXT-XR cultures grown on xylose, and after 12 days for cultures grown on xylooligosaccharides (Table 2). In all cases, nitrogen limitation increased the lipid accumulation, as expected. YBLG3 efficiently transformed cellobiose into lipids, producing up to 20 % of lipids related to total dry weight under nitrogen limitation conditions (Table 2). Performance with glucose as feedstock was slightly lower at both C/N ratios, accumulating 18 % with the highest C/N ratio. YBXT-XR produced almost 35 % of lipids using xylose as substrate in 6 days of culture while achieving lower performance with xylooligosaccharides (14.8 %) after 12 days (Table 2).

**Table 2.** Proportion of lipids over dry biomass (%) accumulated by YBGL3, YBXT-XR and YBXT-XR-BGL3 using glucose (21 g L^-1^) during 3 days, cellobiose (19 g L^-1^ and 1 g L^-1^ glucose) during 3 days, xylose (21 g L^-1^) during 6 days and/or xylooligosaccharides (18 g L^-1^ and 2 g L^-1^ xylose) during 12 days under nitrogen non-limiting (C/N 5) and limiting (C/N 75) conditions.

| Strain | Substrate | C/N 5 | C/N 75 |
| --- | --- | --- | --- |
| YBGL3 | Glucose | 13.5 ± 1.1 | 18.5 ± 1.3 |
|  | Cellobiose | 16.7 ± 2.3 | 19.6 ± 1.0 |
| YBXT-XR | Xylose | 21.1 ± 0.4 | 34.7 ± 0.4 |
|  | Xylooligosaccharides | 9 ± 0.3 | 14.8 ± 0.1 |
| YBXT-XR-BGL3 | Glucose | 20.0 ± 1.6 | 22.9 ± 1.2 |
|  | Cellobiose | 20.2 ± 1.8 | 23.6 ± 1.4 |
|  | Xylose | 27.3 ± 1.8 | 36.4 ± 2.0 |
|  | Xylooligosaccharides | 12.0 ± 0.8 | 16.0 ± 0.6 |

Lipid production was also evaluated for the multifunctional strain in the same conditions described above for the single-enzyme strains (Table 2). Under nitrogen limiting conditions, the yield was similar to those obtained for YBGL3 and YBXT-XR in all the tested substrates (glucose, cellobiose, xylose and xylooligosaccharides), even slightly higher in some cases (Table 2).

### 3.5. Division of labor for lignocellulosic biomass valorization

Division of labor provides a strategy to distribute the metabolic tasks, reducing the burden associated with engineering a single multifunctional chassis. Two strategies were designed to evaluate the contribution of each strain to lignocellulosic biomass monosaccharides processing (Fig. S5). One of them consisted of assessing glucose and xylose consumption in YBGL3 and YBXT-XR monocultures. In the second one, co-cultures of both strains were assembled at different YBGL3:YBXT-XR inoculum ratios (1:1, 1:2, and 1:5), favouring YBXT-XR in order to maximize glucose and xylose co-consumption (Fig. S2). In all cases, independently of the initial inoculum ratio, xylose consumption did not start until glucose was almost completely depleted. At the end of the growth period (114 h), β-xylosidase activity increased with the inoculum ratio, ranging between 20.7 and 28.6 mU mL^-1^, and was 32.5 mU mL^-1^ in the YBXT-XR monoculture. Then, the higher the YBXT-XR inoculum, the higher the final activity.

As β-xylosidase is the limiting activity for lignocellulosic biomass valorization, the three previous ratios and one with higher proportion of YBXT-XR (1:1, 1:2, 1:5 and 1:10) were assayed in co-cultures containing a mixture of 2 g L^-1^ glucose, 8 g L^-1^ cellobiose and 10 g L^-1^ xylooligosaccharides (Fig. 4). In these experiments, increasing the proportion of YBXT-XR resulted in lower cellular densities, since cellobiose cannot be efficiently hydrolyzed when the YBGL3 inoculum is too low. For the 1:10 consortium, where YBGL3 inoculum is 0.025 inoculum), the cellobiose concentration at 144 h was 1.3 g·L□¹. In contrast, complete cellobiose depletion was observed with the 1:1, 1:2, and 1:5 ratios, although the lower the initial YBGL3 fraction, the later the peak of β-glucosidase activity occurred (24, 48, and 72 h, respectively). As a result, cellobiose was exhausted by 48 h with the 1:1 and 1:2 ratios, whereas the 1:5 ratio required 72 h.

**Figure 4.**
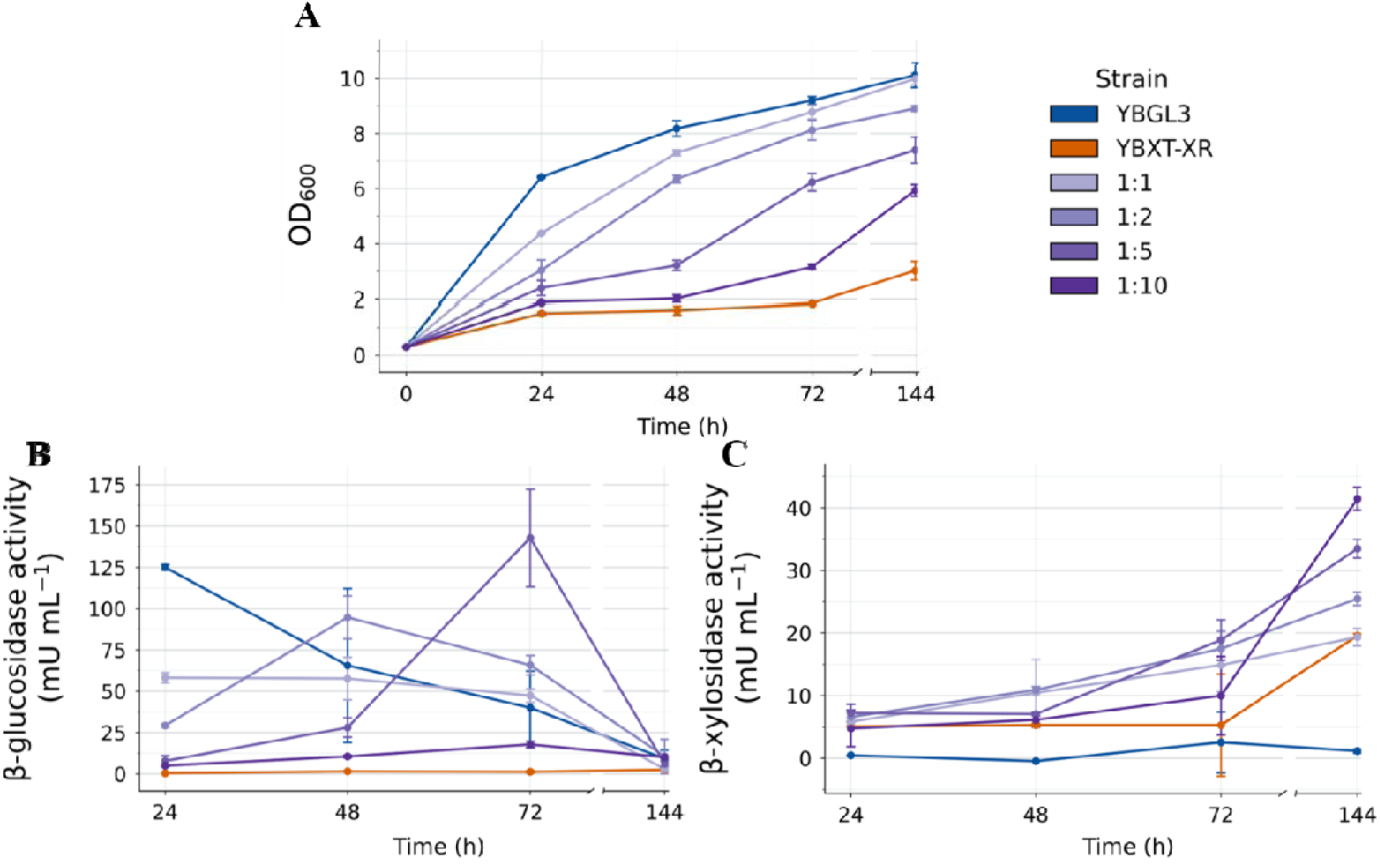
YBGL3 (blue), YBXT-XR (orange) and YBGL3:YBXT-XR consortia (purple) growth on glucose 2 g L^-1^, cellobiose 8 g L^-1^ and xylooligosaccharides 10 g L^-1^. **A)** Optical density at 600nm. **B)** β-glucosidase activity on pNPG. **C)** β-xylosidase activity on pNPX. Error bars correspond to the standard variation of biological triplicates.

β-Xylosidase activity (Fig 4C) displayed a similar trend to that observed in glucose–xylose cultures (Fig. S5). Consortia with higher initial YBXT-XR proportions reached higher final β-xylosidase activities. However, the YBXT-XR monoculture produced the lowest overall growth of tested strains and consortium as it is not able to use the cellobiose (Fig. 4).

### 3.6. Lipid production in consortium vs monoculture

The YBXT-XR-BGL3 monoculture and the YBGL3 + YBXT-XR cocultures were grown in glucose 2 g L^-1^, cellobiose 8 g L^-1^ and xylooligosaccharides 10 g L^-1^ under non-limiting (C/N 5) and nitrogen limiting (C/N 75) conditions. The inoculum ratios selected for the consortium were 1:2 and 1:5 based on previous results. Optical density, enzymatic activity, population ratios and lipid content were followed for 168 h (Fig. 5). The YBXT-XR-BGL3 monoculture reached higher OD600nm than the consortia (Fig. 5A) and secreted higher β-glucosidase (Fig. 5B) and β-xylosidase activities (Fig. 5C), as in the monoculture all the cells were expressing both enzymes. However, in the consortium, the secretion of each enzyme varied depending on the populations’ proportion. HPLC data revealed that, under carbon limiting conditions, cellobiose is completely consumed in 72 h in the consortium and in 48 h in the monoculture [data not shown]. As YBGL3 only can metabolize glucose, YBXT-XR predominated in the consortium after cellobiose depletion, representing nearly 100 % (Table S3) of the culture’s population at 136 h and using xylooligosaccharides to support growth and lipid accumulation. This situation can explain the lower β-glucosidase activity of consortia compared to the YBXT-XR-BGL3 monoculture. By contrast, β-xylosidase activity (Fig. 5C) was similar across all conditions, as almost all cells in the consortium and the monoculture (Table S3) are expressing and secreting BxTw1 from 72 h onwards.

**Figure 5.**
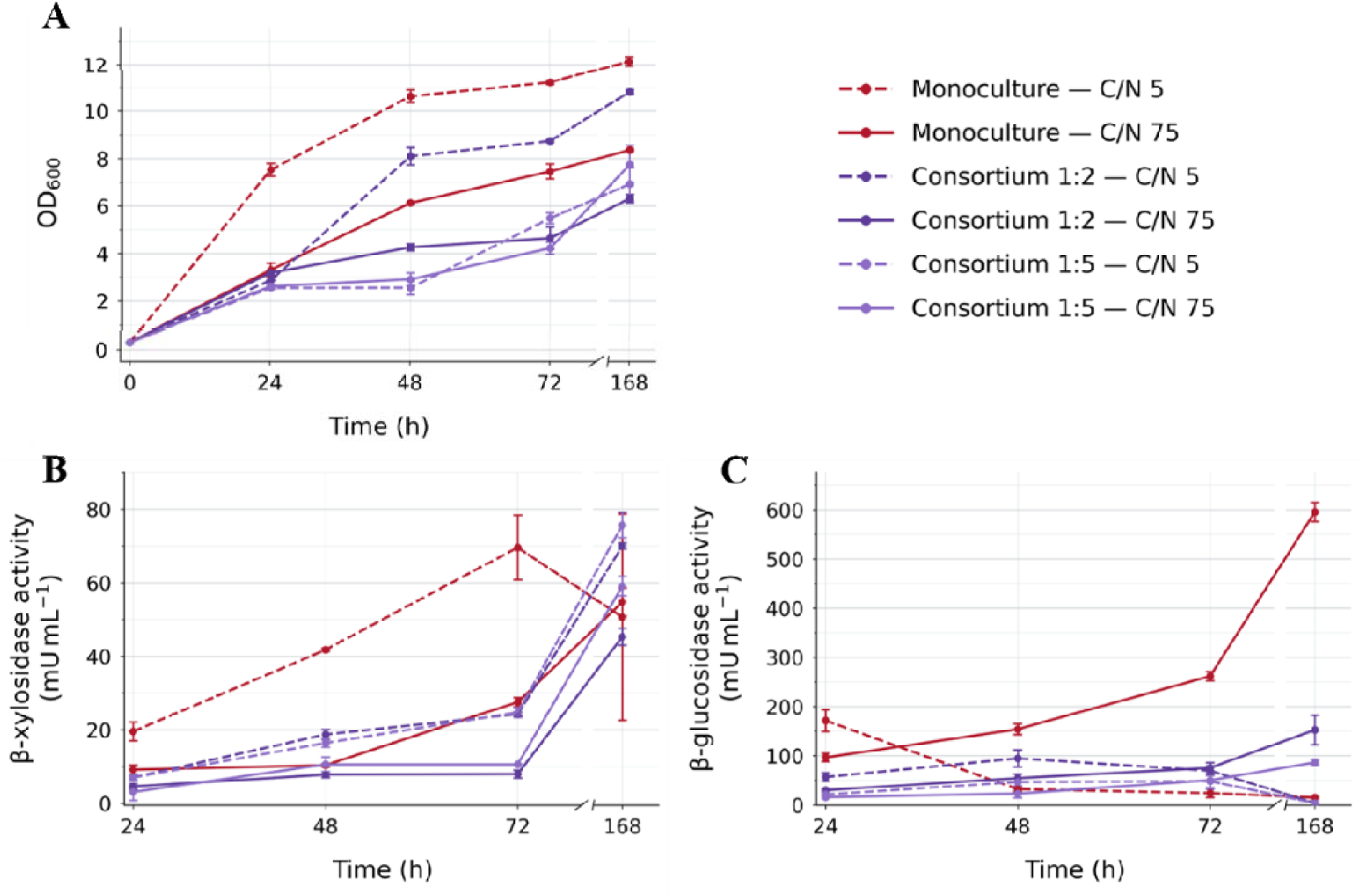
YBXT-XR-BGL3 monoculture (red), consortium 1:2 (dark purple) and consortium 1:5 (light purple) production of lipids on glucose 2 g L^-1^, cellobiose 8 g L^-1^ and xylooligosaccharides 10 g L^-1^ at C/N 5 (dashed lines) or C/N 75 (solid line). **A)** Optical density at 600nm. **B)** β-glucosidase activity on pNPG. C) β-xylosidase activity on pNPX. Error bars correspond to the standard variation of biological duplicates.

Dry biomass production (measured previous to the in the lipid analysis) (Table 3) closely mirrored the optical density profiles observed throughout the experiment (Fig. 5A). The monoculture, where all cells could assimilate both glucose and xylose, yielded more biomass than the consortia. YBGL3 in the consortia enters in starvation conditions after 72 h since it is unable metabolize xylose. OD600nm and biomass production of the consortia grown under nitrogen limitation (C/N 75) were similar, but lipid production with the 1:5 inoculum duplicated that of the ratio 1:2 (0.67 vs. 0.33 g L^-1^ at 168 h) (Table 3). β-xylosidase activity in the former was higher during the whole experiment as initial YBXT-XR inoculum was also higher (Fig. 5C), improving xylooligosaccharaides degradation. YBGL3 represented 30 % and 20 % of the population in consortia inoculated at ratios 1:2 and 1:5, respectively (Table S3), and the performance in lipid productivity is reduced in the condition where starving YBGL3 were more abundant.

**Table 3.** Lipid production (g L^-1^), lipid proportion over dry biomass (%) and biomass production (g L^-1^) of the consortium and monoculture under nitrogen non-limited (C/N 5) or limited (C/N 75) conditions.

| Culture | C/N ratio | Time (h) | Lipid conc. (g L<sup>-1</sup>) | Lipid prop. (%) | Biomass conc. (g L<sup>-1</sup>) |
| --- | --- | --- | --- | --- | --- |
| Consortium 1:2 | 5 | 72 | 0.26 ± 0.00 | 8.7 ± 0.1 | 3.04 ± NA |
|  |  | 120 | 0.39 ± 0.06 | 12.7 ± 1.9 | 3.26 ± NA |
|  |  | 168 | 0.34 ± 0.02 | 11.1 ± 0.6 | 3.29 ± NA |
|  | 75 | 72 | 0.23 ± 0.00 | 12.9 ± 0.0 | 1.82 ± NA |
|  |  | 120 | 0.30 ± 0.00 | 16.5 ± 0.0 | 1.73 ± NA |
|  |  | 168 | 0.33 ± 0.01 | 18.3 ± 0.6 | 1.86 ± NA |
| Consortium 1:5 | 5 | 72 | 0.23 ± 0.01 | 10.6 ± 1.8 | 2.18 ± 0.07 |
|  |  | 120 | 0.30 ± 0.00 | 11.5 ± 3.4 | 2.63 ± 0.04 |
|  |  | 168 | 0.40 ± 0.02 | 11.1 ± 0.6 | 2.59 ± 0.16 |
|  | 75 | 72 | 0.23 ± 0.00 | 15.3 ± 0.4 | 1.49 ± 0.02 |
|  |  | 120 | 0.37 ± 0.00 | 19.2 ± 1.2 | 1.91 ± 0.01 |
|  |  | 168 | 0.67 ± 0.01 | 34.0 ± 2.3 | 1.96 ± 0.02 |
| Monoculture | 5 | 72 | $0.54 \pm 0.00$ | $13.1 \pm 0.0$ | $4.16 \pm \text{NA}$ |
| | | 120 | $0.64 \pm 0.01$ | $15.4 \pm 0.2$ | $3.94 \pm \text{NA}$ |
| | | 168 | $0.58 \pm 0.03$ | $14.1 \pm 0.8$ | $3.94 \pm \text{NA}$ |
| | 75 | 72 | $0.46 \pm 0.03$ | $18.4 \pm 1.1$ | $2.50 \pm \text{NA}$ |
| | | 120 | $0.63 \pm 0.03$ | $25.3 \pm 1.1$ | $2.56 \pm \text{NA}$ |
| | | 168 | $0.66 \pm 0.04$ | $26.3 \pm 1.5$ | $2.34 \pm \text{NA}$ |

Accordingly, differences in biomass formation and population dynamics were reflected in the temporal patterns of lipid accumulation. The monoculture showed the highest lipid at 72 and 120 h, however, the 1:5 consortium ultimately reached a comparable lipid titer at 168 h. This was driven by a pronounced increase in lipid content during the final 48 h, from 19.2 % to 34.0 % during the final 48 h, as well as lipid production, accompanied by an increase in lipid production from 0.37 to 0.67 g L□¹. During this phase, biomass no longer increased, indicating that carbon was almost exclusively redirected toward lipid synthesis. Between 72 and 168 h IN THE YBXT-XR-BGL3 monoculture, no substantial change in lipid content or biomass was detected as the carbon source was probably fully consumed (no xylooligosaccharides between 2 and 5 xylose units were detected). With respect to total lipid productivity, the 0.66 ± 0.04 g L^-1^ at 120 h and the 1:5 consortium 0.67 ± 0.01 g L^-1^.

## 4. Discussion

This study shows how *Y. lipolytica* can be engineered to depolymerize and metabolize both cellulose- and hemicellulose-derived oligosaccharides, and that a division-of-labor strategy provides a robust framework to distribute these metabolic tasks across specialized strains. By combining heterologous secretion of *T. amestolkiae* β-glucosidase (BGL3) and β-xylosidase (BxTw1) with the xylose reductase pathway, we established a multifunctional monoculture and a two-member consortium capable of processing the main polysaccharide fractions of lignocellulosic biomass (LCB). These systems provide insight into the trade-offs between enzymes’ secretion metabolic burden, substrate specialization, and cooperative processing, which are critical factors for developing efficient bioprocesses for LCB valorization.

### 4.1. Development of a multifunctional Y. lipolytica strain for lipid production from lignocellulose-derived oligosaccharides

*Y. lipolytica* was able to successfully express and secrete functional fungal glycosidases despite the intrinsic challenges associated with heterologous expression and secretion of fungal enzymes. Engineering a multifunctional strain capable of both degrading cellulose- and hemicellulose-derived substrates and metabolizing xylose is often limited by metabolic burden, especially when the secretory pathway and cofactor-imbalanced sugar catabolism are simultaneously expressed ^18^. However, YBXT-XR-BGL3 activity on each substrate was comparable to those of the individual strains bearing expressing BGL3 or BxTw (Figs. 1 and 3). This robustness contrasts with previous studies showing that co-expression of multiple cellulolytic enzymes in *Y. lipolytica* significantly reduced growth and lipid accumulation unless mitigated by chaperone expression ^33^ or by adding easily metabolizable carbon sources ^31^. In our case, the high catalytic efficiency of BGL3 at pH 5 and 6 was sufficient to ensure rapid cellobiose hydrolysis and assimilation in the multifunctional and YBGL3 strains.

Lipid production in monocultures was assayed at pH 6 to balance enzyme performance with cellular stability. Although saccharification yields at pH 5 were slightly higher, *Y. lipolytica* typically acidifies its environment during growth ^55^, and operating at a higher starting pH prevents excessive acidification at late fermentation stages. Under these conditions, YBXT-XR-BGL3 accumulated ∼20 % lipids from cellobiose, like YBGL3. This amount was slightly higher than the production from glucose, highlighting the efficiency of engineering *Y. lipolytica* for lipid production from cellobiose. Previous studies reported on a strain capable of metabolizing cellobiose and cellotriose throughout the expression of a cellobiose phospholyase and a cellodextrin transporter, achieving growth rate and biomass comparable to those obtained on glucose ^56^. Other study ^57^, reported a lipid concentration of 0.8 g L^-1^ from 50 g L^-1^ of microcrystalline cellulose (Avicel) by overexpression of two endogenous, B-glucosidases in *Y. lipolytica*, and and simultaneous Celluclast saccharification, which represent lower carbon to lipid conversion compared to YBXT-XR-BGL3 (0.67 g L^-1^ from 20 g L^-1^ of cellobiose). These results highlight that the degradation cellooligosaccharides can be successfully engineered in *Y. lipolytica*.

Although several studies have reported the use of *Y. lipolytica* to valorize hemicellulose-derived products, into lipids or other compounds such as coumaric acid ^58^, the effective valorization of this material remains challenging. Some studies showed poor growth on xylans of engineered strains due to inefficient xylose release or insufficient intrinsic xylose metabolism ^30,59^. In our case, the strain YBXT-XR-BL3 showed high efficiency accumulating ∼15 % lipids from xylooligosaccharides and around 35 % from xylose. This strain is the first strain to valorise simultaneously cellulose and hemicellulose-derived substrates.

### 4.2. Division of labor impact on lipid production from lignocellulose oligosaccharides in *Y. lipolytica*

Division of labor provides a strategic framework for distributing the metabolic requirements of LCB valorization across multiple specialized strains ^29^. In this context, separating cellulolytic and hemicellulolytic functions between YBGL3 and YBXT-XR offers a way to coordinate depolymerization and assimilation. Glucose and xylose are the main monomers liberated from cellulose and hemicellulose depolymerization, respectively ^60^. It is well-known that even in engineered *Y. lipolytica* for xylose consumption, glucose is the preferred monomer.

We first evaluated whether altering initial population ratios could influence glucose–xylose consumption dynamics. As expected, glucose was consumed first across all conditions, reflecting strong glucose preference and inhibition of xylose uptake ^61^. Once glucose is depleted from the medium, xylose is consumed, as xylose transporters are competitively inhibited by glucose, preventing co-consumption ^61^. This effect has been observed in many organisms across the tree of life ^62^. Nevertheless, initial inoculum composition strongly affected the enzymatic landscape: higher proportions of YBXT-XR led to greater β-xylosidase activity, which is the rate-limiting step for xylooligosaccharides utilization. This indicates that division-of-labor in consortia can be tuned by adjusting the initial population ratios of the individual members, even when monosaccharide uptake kinetics is constrained by transport physiology.

YBGL3 rapidly hydrolyzed cellobiose, releasing glucose that fuelled early exponential growth and supported BxTw1 secretion by YBXT-XR. If YBGL3 was underrepresented glucose release was delayed or incomplete, limiting overall performance. Conversely, high initial YBXT-XR proportions ensured sustained β-xylosidase activity later in the process, once cellobiose had been depleted and xylooligosaccharides became the dominant carbon source.

These results highlight the mechanistic advantage of division of labor: each strain specializes in a distinct task and contributes optimally at different stages of substrate turnover, enabling dynamic distribution of metabolic roles as the available carbon pool evolves. Properly tuning inoculum ratios is therefore critical for maximizing cooperative processing of mixed lignocellulosic substrates.

Lipid production under nitrogen-limiting conditions provided an accurate metric for comparing the functional efficiency of consortia with that of monocultures. The 1:5 consortium achieved lipid titers comparable to those of the engineered YBXT-XR-BGL3 monoculture, whereas the 1:2 consortium displayed markedly lower production (Table 3). This divergence can be rationalized by considering population dynamics, carbon allocation and nitrogen concentration. In the 1:5 ratio, the high initial abundance of YBXT-XR supported strong β-xylosidase activity and xylose availability throughout fermentation allowing this strain to keep on growing (Fig. 5). However, quick cellobiose degradation and glucose consumption by both strains caused carbon starvation of YBGL3. In contrast, in cultures inoculated with the 1:2 ratio, the YBGL3 subpopulation experienced prolonged starvation once glucose was depleted (Table S3), leading to lipid mobilization ^63^ and reducing culture-level yields. The multifunctional monoculture YBXT-XR-BGL3 reached lipid titers comparable to those of the best-performing consortium, confirming that co-expression of BGL3, BxTw1, and the xylose pathway does not impose a prohibitive metabolic load under the tested conditions.

In the consortium, YBGL3 and YBXT-XR are competing for glucose uptake and metabolism and once it is depleted, YBXT-XR starts using xylose. To reduce the level of competition and increase consortium performance, it would be desirable to knock out of glucose metabolism in YBXT-XR strain or engineer real cross feeding by developing a consortium between a YBGL3-XR strain unable to consume glucose and YBXT ^38^. However, even in this scenario, glucose would be uptake first than xylose due to transporters affinity. Engineering of sugar transporters that are able of transporting xylose in the presence of glucose ^64–66^ and its expression in *Y. lipolytica* to achieve true co-consumption of both sugars would be highly desirable for the development of cost-effective bioprocess ^67^.

## 5. Conclusions

In this work, we demonstrate that *Y. lipolytica* can be engineered to efficiently depolymerize and metabolize both cellulose- and hemicellulose-derived oligosaccharides through the secretion of *T. amestolkiae* BGL3 and BxTw1 and the integration of a functional xylose reductase pathway. We show that these activities can be combined either within a single multifunctional chassis or, more strategically, distributed across a two-strain consortium following a division-of-labor approach. Both architectures supported lipid production from mixed lignocellulosic substrates, with the monoculture displaying robust performance and the consortium offering tuneable specialization through inoculum ratio adjustment. Although co-consumption of glucose and xylose remained constrained by native transport physiology, our results reveal clear opportunities for further optimization, such as engineering cross-feeding interactions or glucose-insensitive xylose transport, to unlock the full potential of cooperative systems. Collectively, this study establishes *Y. lipolytica* as a versatile platform for consolidated bioprocessing and highlights division of labor as a promising strategy to advance the efficient and modular valorization of lignocellulosic biomass.

## Acknowledgements

This work was supported by the Spanish projects MICODE (PID2020-114210RB-I00 MCIN/AEI) and MOLA (PID2024-162673NB-I00 MCIN/AEI), and the European projects FRONTSH1P (H2020 101037031). The authors acknowledge the support toward the publication fee by the CSIC Open Access Publication Support Initiative through its Unit of Information Resources for Research (URICI).

## SUPPLEMENTARY MATERIAL

**Figure S1.**
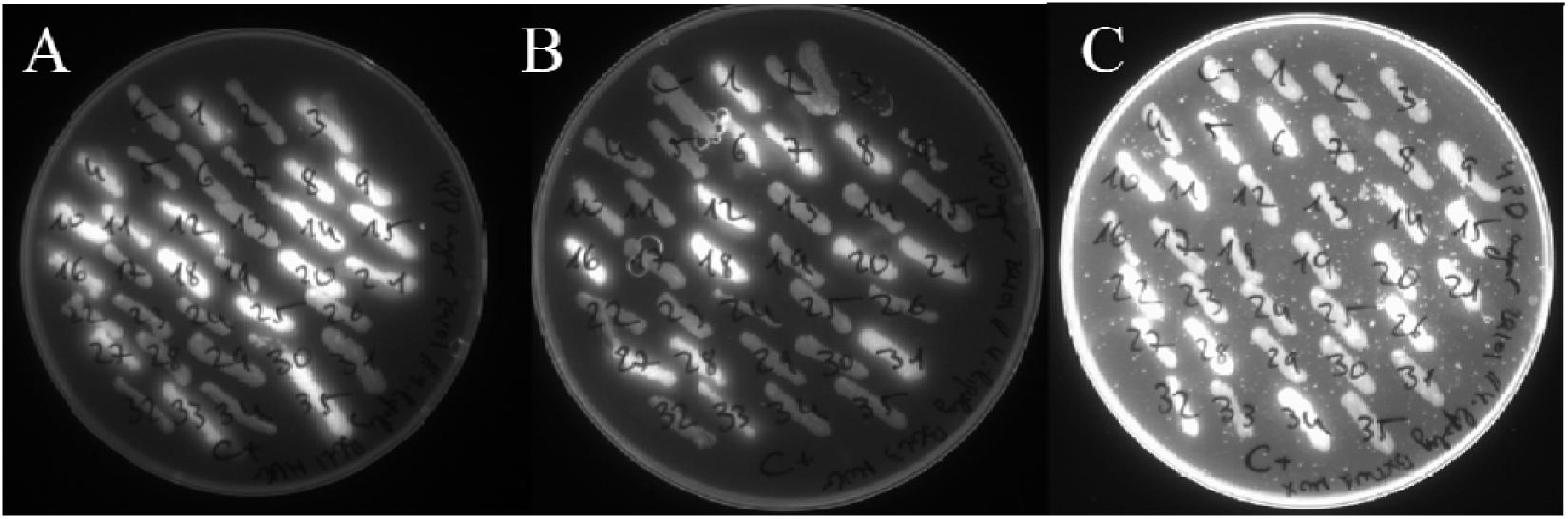
Fluorescence of transformed candidates from YBGL1 (*A*), YBGL3 (*B*) and YBXT (*C*) after incubation with 4-methylumbelliferyl-β-D-glucopyranoside (*A* and *B*) or 4-methylumbelliferyl-β-D-xylopyranoside (*C*). Negative control corresponds to the first colony at the upper left part.

**Figure S2.**
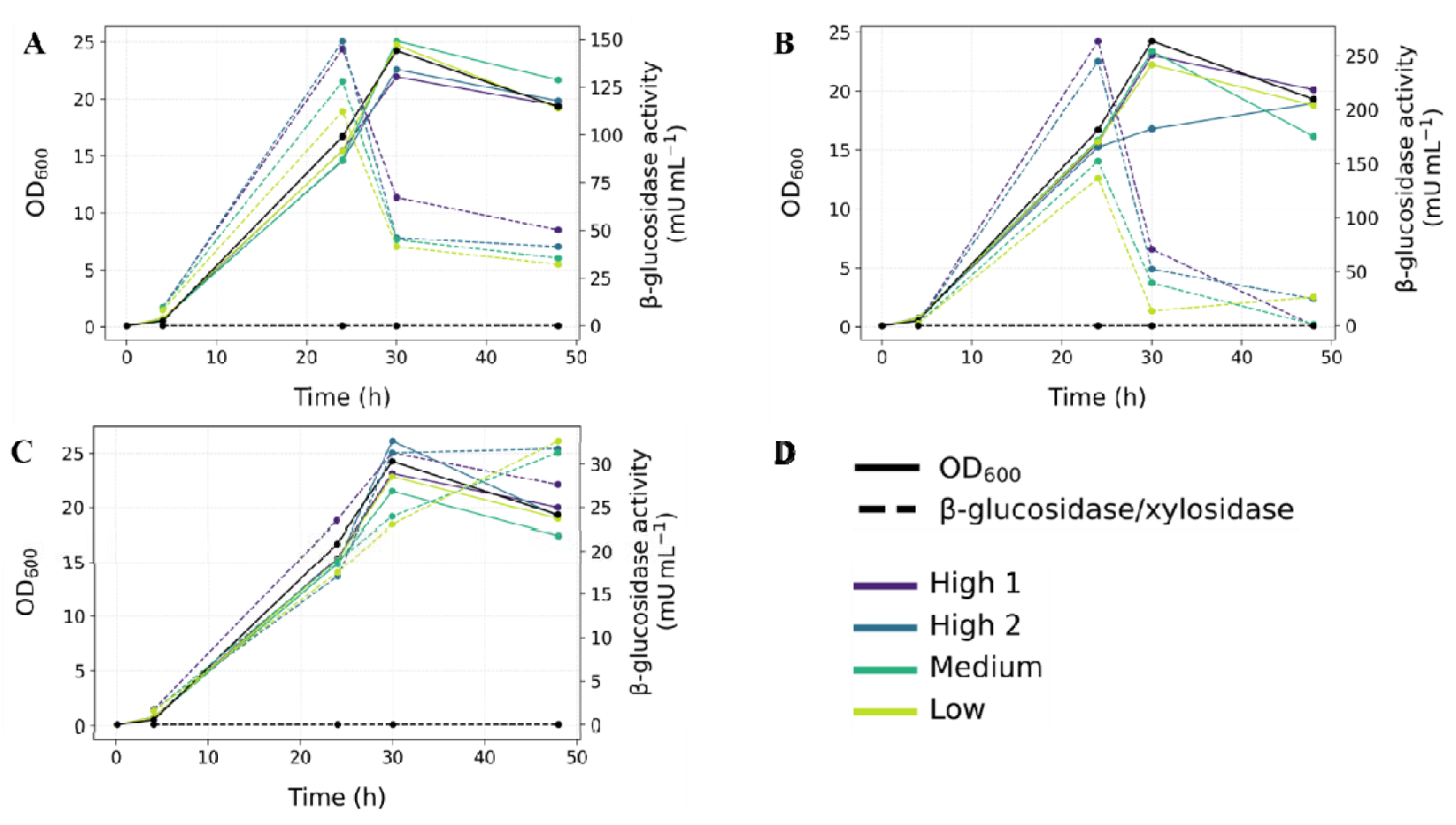
Selected clones from YBGL1 (*A*), YBGL3 (*B*) and YBXT (*C*) transformation screening. Optical density (OD_600nm_) and enzymatic activities (β-glucosidase for YBGL1 and YBGL3, or β-xylosidase for YBXT) are displayed in solid and dashed lines respectively. Tested clones are High1 (purple), High2 (blue), Medium (green) and Low (yellow) depending on the fluorescence displayed in the MUG/MUX assay.

**Figure S3.**
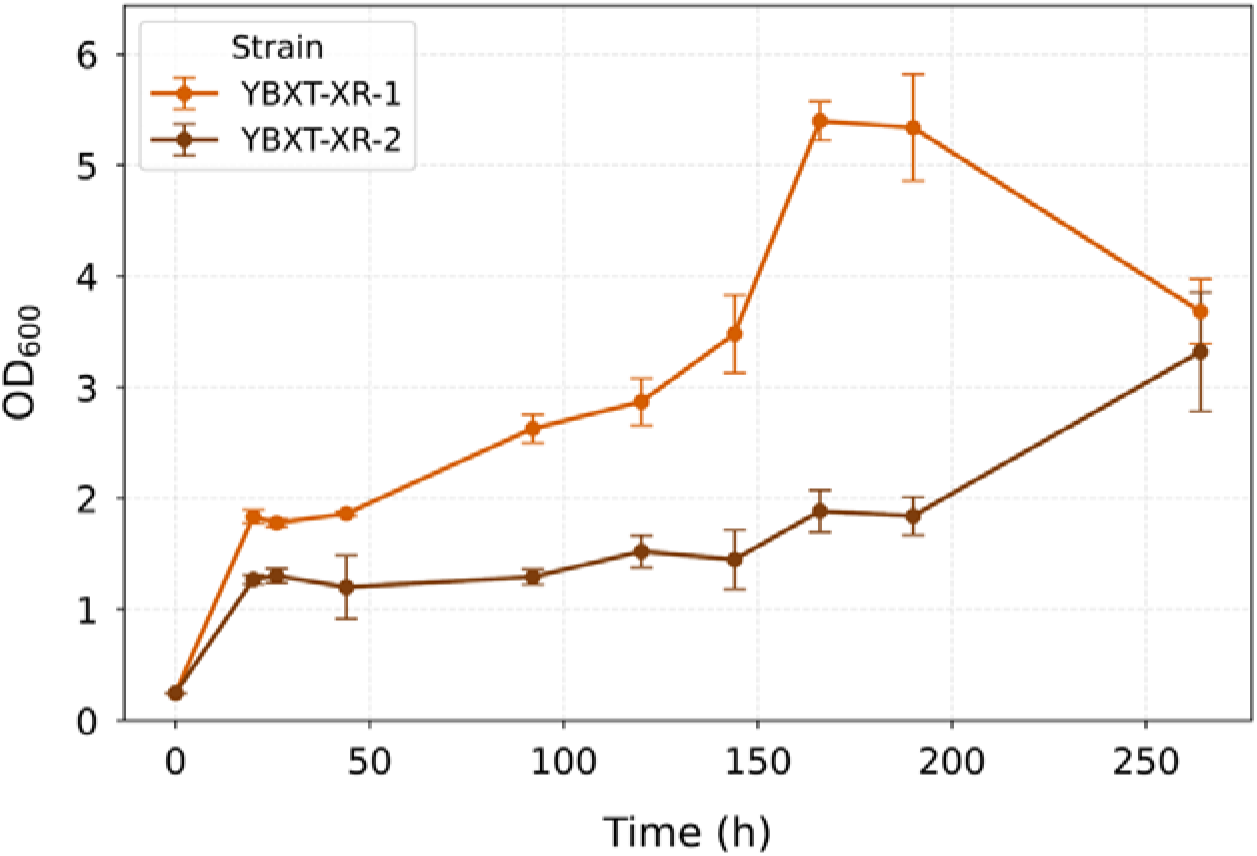
Optical density (OD_600nm_) of two YBXT-XR-BGL3 transformants (transformant 1, orange; transformant 2, brown) on YNB with 10 g L^-1^ xylooligosaccharides and 1 g L^-1^ xylose at pH 6.

**Figure S4.**
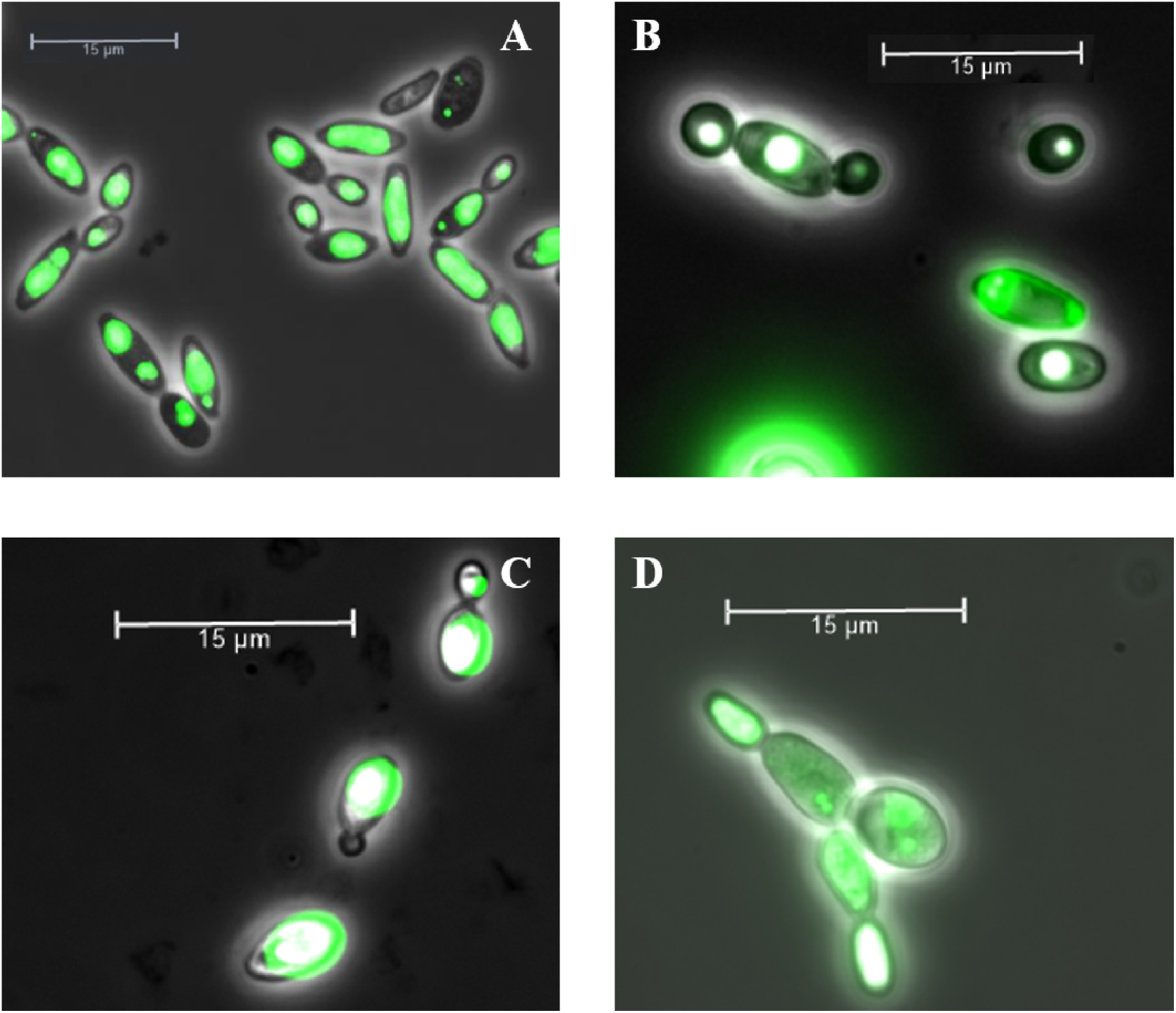
Fluorescence microscopy pictures of YBGL3 strain on cellobiose 19 g L^-1^ and glucose 1 g L^-1^ (*A*), YBXT strain on xylooligosacccharides 19 g L^-1^ and xylose 1 g L^-1^ (*B*), YBXT-XR-BGL3 strain on cellobiose 19 g L^-1^ and glucose 1 g L^-1^ (*C*) and xylooligosacccharides 19 g L^-1^ and xylose 1 g L^-1^ (*D*) under carbon limiting conditions (C/N 75). Cells were stained with BODIPY and visualized by fluorescence microscopy. Scale bars indicate 15μm.

**Figure S5.**
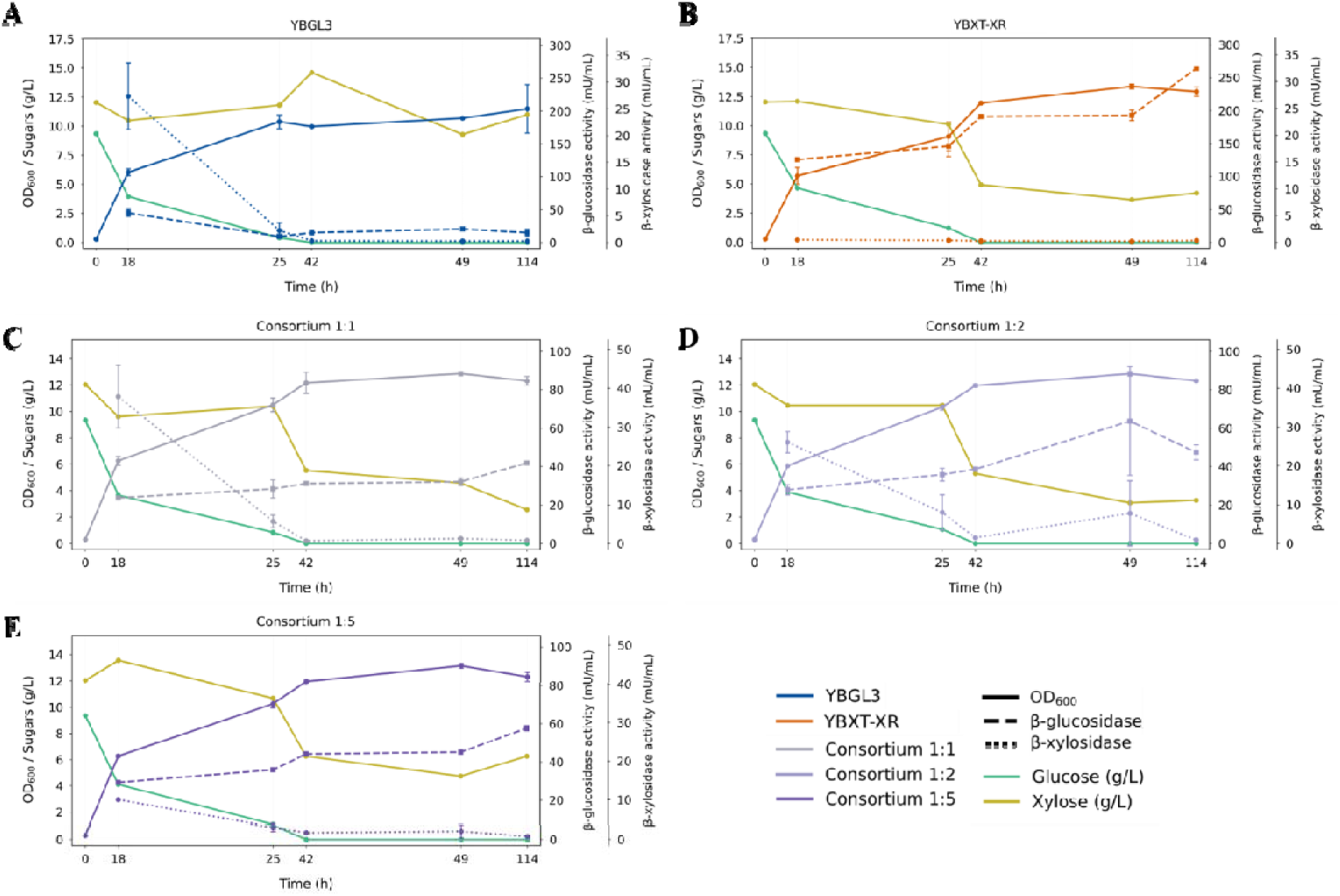
YBGL3 monoculture (*A*, blue), YBXT-XR monoculture (*B*, orange), YBGL3:YBXT-XR consortium 1:1 (*C*, grey), 1:2 (*D*, light purple) and 1:5 (*E*, purple) growth over glucose 10 g L^-1^ and xylose 10 g L^-1^. Optical density (OD_600nm_, solid line), β-glucosidase activity (dashed line), β-xylosidase activity (dotted line), glucose (green) and xylose (yellow) concentration are represented. Error bars correspond to the standard variation of biological triplicates.

**Table S1.**
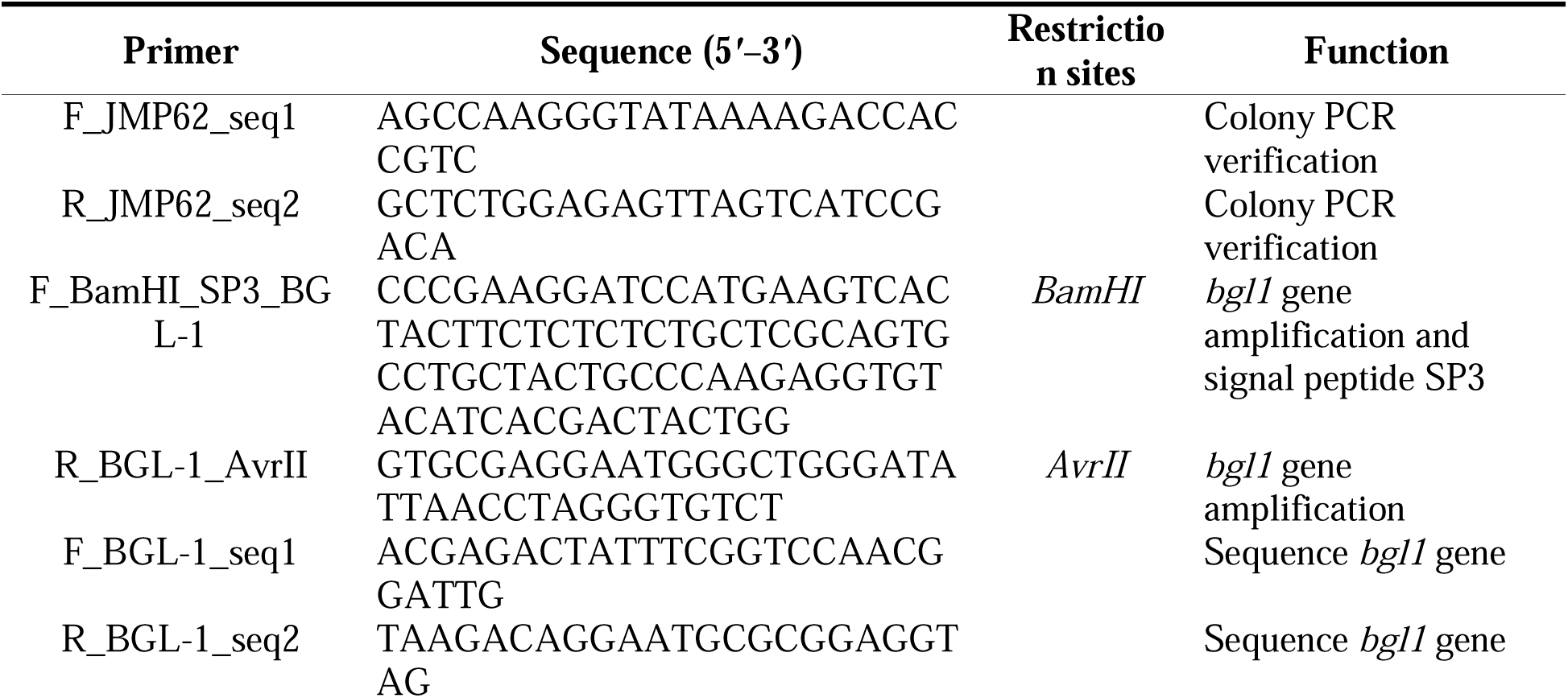
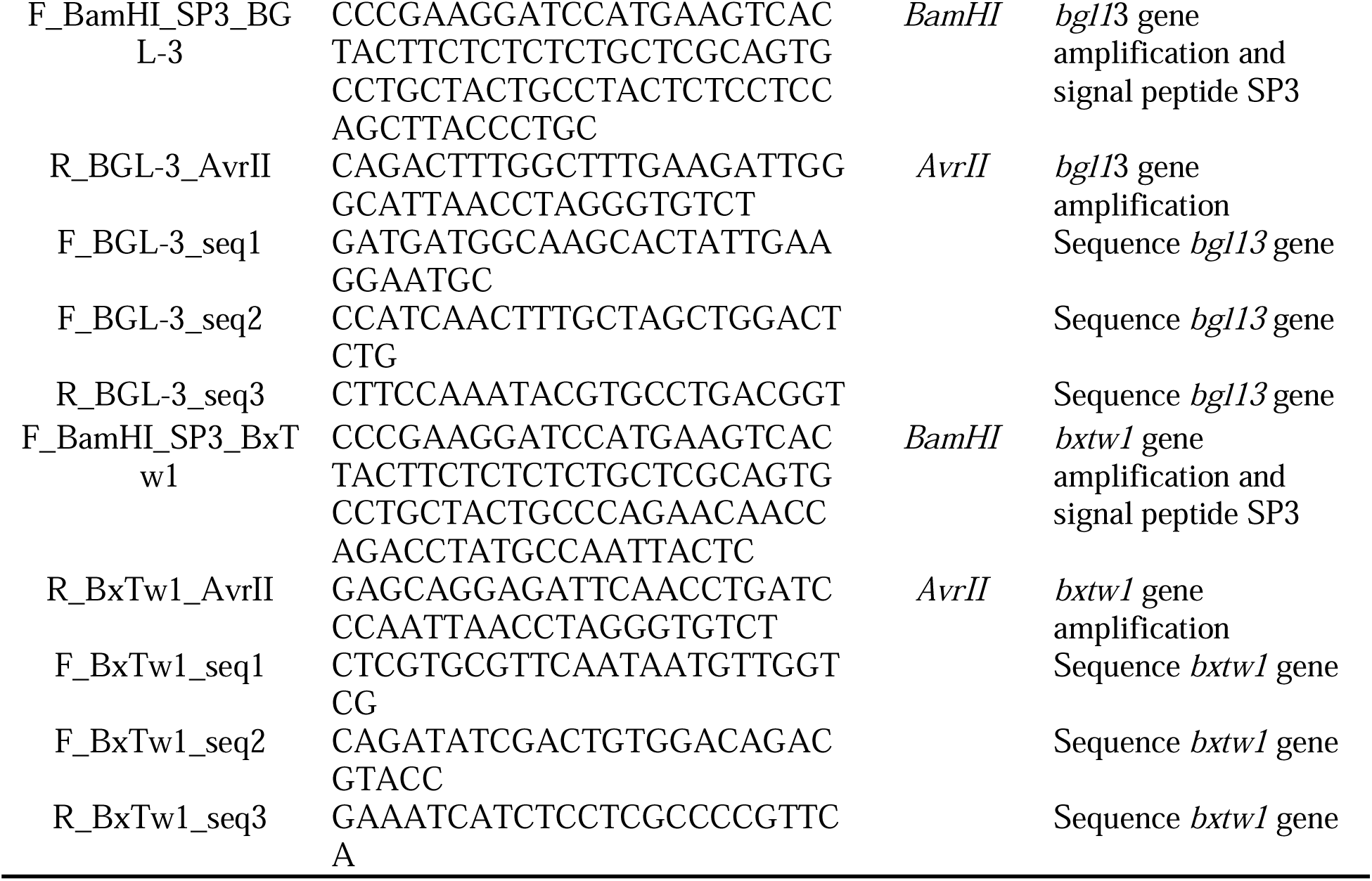
Primers used in this study.

**Table S2.**
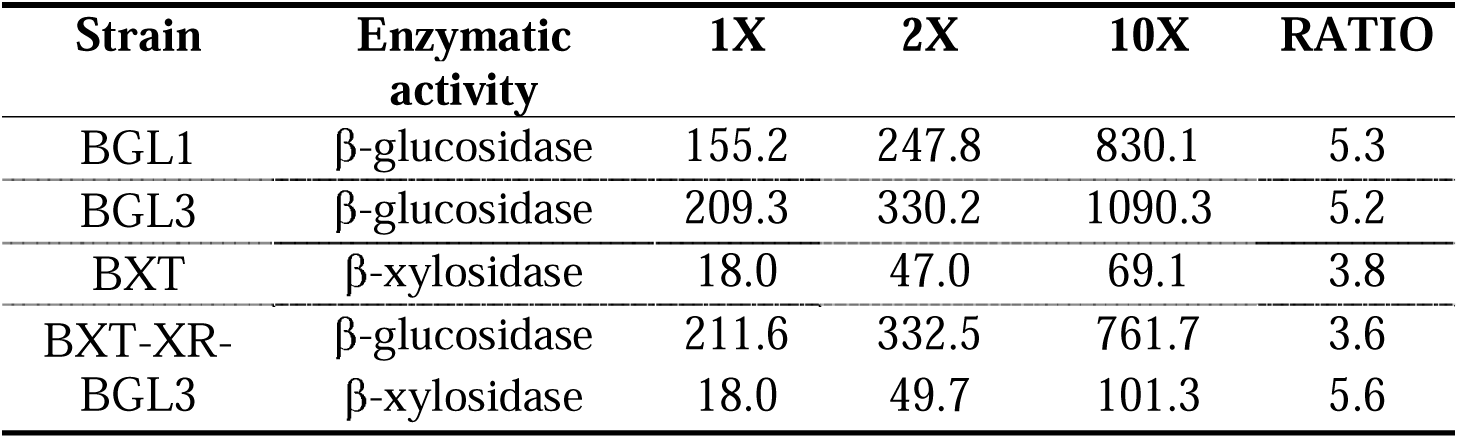
Supernatant concentration of the strains. The initial volume was 50 mL. **1×** indicates enzymatic activity measured without concentration, **2×** indicates a twofold concentration (final volume 25 mL), and **10×** indicates a tenfold concentration (final volume 5 mL).

**Table S3.** Cellular proportions in the consortium represented as the percentage (%) of YBXT-XR cells in the culture at different sampling points.

| Culture | C/N ratio | 24 h | 72 h | 136 h |
| --- | --- | --- | --- | --- |
| Consortium 1:2 | 5 | 70 | 93 | 100 |
|  | 75 | 84 | 70 | 100 |
| Consortium 1:5 | 5 | 96 | 90 | 97 |
|  | 75 | 93 | 80 | 100 |

## Notes

### Competing Interest Statement

The authors have declared no competing interest.

